# Transcriptomic Enrichment of Ferroptosis-Related Gene *ACSL4* in Advanced Hepatic Fibrosis/Cirrhosis: Bioinformatics Analysis and Experimental validation

**DOI:** 10.1101/2023.06.22.543838

**Authors:** Shuya Zhang, Ying Liu, Liping Chen, Yuxuan Liu, Yuqi Guo, Jilin Cheng, Jun Huang

## Abstract

**Background:** Liver fibrosis is a critical part of the clinical process of liver disease that progresses to cirrhosis and even liver cancer, and effective treatment and clinical biomarkers are urgently needed to manage liver fibrosis. Ferroptosis, a notable biological phenomenon that has received attention because of the role it performs in liver fibrosis. The objective of this research is in order to identify key ferroptosis genes related to advanced liver fibrosis/cirrhosis.

**Methods:** Gene expression differences were analyzed in liver fibrosis liver tissue of hepatitis B virus(HBV)infection patients, non-alcoholic steatohepatitis (NASH) patients and alcoholic hepatitis (AH) patients to obtain overlapping ferroptosis-related genes that are significantly up-regulated. A multigroup comparison was performed to evaluate the diagnostic clinical importance of ferroptosis-related genes of patients in differential degrees of liver fibrosis, and confirmed via gene expression trend analysis.

The differential expression of candidate ferroptosis-related genes through classical carbon tetrachloride (CCl_4_) induced advanced liver fibrosis mice model were validated by real-time quantitative PCR (qPCR). Correlation analysis was conducted to tentatively identify the connections between hepatic ferroptosis-related genes and key genes participating in functional pathways relevant to liver fibrosis.

**Results:** We screened and obtained 10 genes related to ferroptosis, all of which were significantly up-regulated in liver tissue from liver fibrosis patients of different etiologies, and identified acyl-CoA synthetase long chain family member 4 (*ACSL4*) was transcriptomic enriched in patients with HBV infection, NASH, AH-associated advanced liver fibrosis and cirrhotic tissue adjacent to hepatocellular carcinoma (HCC). In CCl_4_ induced advanced liver fibrosis mice model, the hepatic *ACSL4* expression was significantly up-regulated when compared to normal controls. In our study, we also suggest a significant association between *ACSL4* and representative genes in liver fibrosis-related pathway.

**Conclusion:** We found that *ACSL4* mRNA can effectively differentiate the severity of liver fibrosis, suggesting its potential clinical diagnostic value in patients with liver fibrosis regardless of its etiology. *ACSL4* may be a promising biomarker, which deserves further research.

## Introduction

Chronic liver disease is the primary cause of the global health burden, which is one of the world’s significant public health issues.^[1]^. Based on statistical data, more than 840 million people worldwide currently suffer from different types for chronic liver disease, causing about 2 million deaths each year^[2-4]^. Liver fibrosis plays the necessary role in the advancement of chronic liver disease, which has been considered to be one of the reactions to healing liver damage, and its ongoing progression without intervention can cause hypertension and hepatic encephalopathy, with other clinical complications^[5-8]^. If patients with severe liver fibrosis are not treated promptly and effectively, the condition can progress to irreversible cirrhosis or even result in functional liver failure^[9-11]^°

In general, liver fibrosis is initially induced by liver injury due to different types of causative agents, which include viruses (hepatitis B virus and hepatitis C virus) ^[12-14]^, alcoholism (alcoholic steatohepatitis)^[15, 16]^, non-alcoholic factors (non-alcoholic steatohepatitis)^[17, 18]^, and some immune factors (autoimmune diseases and genetic diseases)^[19]^. Liver fibrosis is a dynamic, multilevel molecular, cellular and tissue co-activation process. The extracellular matrix (ECM) component gradually over-accumulates during the disease process and is maintained by the continuous activation of hepatic myofibroblasts^[20]^.Current studies consistently indicate that regardless of its etiology, the onset and progression of liver fibrosis follows a series of common events^[21]^. These events include the release of dangerous or death-related molecular patterns and the hepatic stellate cells (HSCs) activation and then conversion to myofibroblasts, resulting in the excessive accumulation of ECM. Progressive ECM deposition leads to the destruction of the liver’s structural integrity, including the loss of hepatocytes, ultimately culminating in the formation of pseudobulbar structures, which further progress to cirrhosis. Ultimately, advanced liver fibrosis results in fatal liver dysfunction and liver failure^[22]^.

Ferroptosis represents a recently discovered novel form of programmed cell death mainly through glutathione peroxidase 4 (*GPX4*) inactivation^[23]^. Ferroptosis accompanies unique biological features, including intracellular iron overload and lipid peroxidation, ROS accumulation^[24]^, cell swelling, abnormal mitochondrial morphology, and cell lysis^[25]^. It was shown that excessive hepatic iron deposition and ferroptosis enhance acetaminophen induced liver fibrosis in mice^[26]^. Ferroptosis regulates the autophagic signaling pathway in HSCs^[27]^. Long-chain acyl coenzyme A synthase 4 (*ACSL4*) is a critical gene involved in ferroptosis ^[28]^ which is related to several metabolic diseases ^[29-31]^, and it has been shown that hepatic *ACSL4* in patients with NAFLD exhibits high levels of expression^[32]^, and inhibition of *ACSL4* ameliorates liver fibrosis caused of NASH in a mice model ^[33]^. However, there is currently no research on the expression of hepatic *ACSL4* in the staging of liver fibrosis induced by differential etiologies, and whether it is dysregulated at specific stages remains uncertain.

In this study, we initially discovered that *ACSL4* is a gene that exhibits upregulation in liver fibrosis cases related to HBV infection, NASH, and AH. Moreover, when considering liver fibrosis severity, the gene was specifically upregulated in liver tissue in advanced fibrosis/cirrhosis. Subsequently, the expression levels of *Acsl4* were verified in CCl_4_-induced advanced liver fibrosis mice. Finally, for functional analysis, the correlations between *Acsl4* and hub genes involved in the liver fibrosis pathway were determined.

## Materials and Methods

### Publicly available data

The datasets and clinical information of patients required were downloaded from the Gene Expression Omnibus (GEO) gene-expression data base using the R Studio’s “GEO query” R package (v.4.2.3), and patients with incomplete clinical information were excluded. Four liver tissue datasets GSE84044, GSE48452, GSE103580 and GSE14520 of patients in liver fibrosis/ cirrhosis with control were included. Three liver tissue datasets from classical CCl_4_-induced liver fibrosis mice model and control liver tissue including GSE55747, GSE130123 and GSE152250 were involved in this study. Detailed information can be found in TABLE 1.

### Acquisition of differentially up-regulated genes

Differentially up-regulated genes were screened using the “limma” R package for P<0.05 and FC≥1.5 in the disease samples compared to the control samples. The distribution of differentially up-regulated genes between disease and control samples using the “ggplot2” R package, and volcano-like plots of differentially expressed genes were drawn.

### Protein-protein interaction analysis and functional enrichment analysis of differentially upregulated genes

The String database (http://string.embl.de/) was used for Protein-Protein-Interaction (PPI) network map visualization. The “clusterProfiler” R package is used for genomic GO and KEGG functional enrichment analysis, and after specifying the signaling pathways and biological functions involved in commonly up-regulated genes, the “ggplot2 “ R package visualizes the results of enrichment analysis.

### Acquisition of ferroptosis -associated commonly upregulated genes

The commonly upregulated genes from the three diseases were analyzed. Ferroptosis -associated genes in commonly upregulated genes were further screened using Ferrdb Database (http://www.zhounan.org/ferrdb/current/operations/download.html)^[34]^ and by conducting a comprehensive literature review (as of January 2023).

### Other bioinformatics analysis methods

The “Tcseq” R package is used for gene expression time trend analysis. The “cor” package is used for analysis of Spearman’s correlation coefficient, and P<0.05 is considered as statistically significant.

### Animal experiment

Healthy C57BL/6 male mice of 18-20 g weight were selected and fed for one week after acclimatization and divided into control (CTRL) and CCl_4_ groups. Intraperitoneal injections were administered (twice/one week) in the dosage of 1 ml/kg per week for 8 weeks. 25% CCl_4_ olive oil solution was given to the CCl4 group, and the same dosage of olive oil was given to the control group. The animal experiment in this study was approved by Ethics Committee of Zhengzhou University.

### Hematoxylin and Eosin and Sirius red staining

Section of liver tissue were prepared and hematoxylin and eosin (H&E) staining was performed using 3-_μ_m liver tissue sections, and 4-_μ_m liver tissue sections were used for Sirius red staining. The specific steps were formalin fixation, dehydration, paraffin embedding and sectioning of liver tissue, deparaffinization of the sections to water, staining, and sealing sections with neutral gum.

### QPCR detection of *ACSL4* expression

Extract total RNA from liver tissue and reverse transcribe it into cDNA. SYBR green reagent was used for qPCR analysis. *Gapdh* served as an endogenous control. Primers used in the study include the following:

*Gapdh*- forward primer: GCACAGTCAAGGCCGAGAAT;

*Gapdh*- reverse primer: GCCTTCTCCATGGTGGTGAA;

*Acsl4*- forward primer: CCACACTTATGGCCGCTGTT;

*Acsl4*- reverse primer: GGGCGTCATAGCCTTTCTTG.

## Statistical method

Recommended method in R studio (v.4.2.3) was used for statistics analysis according to the characteristics of the GEO database. For the experimental data of the mice model, the statistical analysis was performed in GraphPad Prism 8.0 by t-test. p<0.05 was statistically significant.

## Results

### Transcriptome profiling in patients with HBV, NASH and AH-associated liver fibrosis reveals a total of 25 commonly up-regulated genes

To acquire the common liver fibrosis causative genes, we explored the gene expression characteristics of liver fibrosis associated with different disease etiologies. Firstly, three gene microarray datasets of liver tissue fibrosis caused by HBV, NASH and AH were screened from the GEO public database: the GSE84044 dataset of HBV infection-associated liver fibrosis patients, the GSE48452 dataset of NASH-associated liver fibrosis patients, and the GSE103580 dataset of AH-associated liver fibrosis/cirrhosis patients. More than 200 genes were significantly up-regulated and more than 30 genes were significantly down-regulated in HBV infection-associated liver fibrosis compared to control (Fig. 1A), and more than 300 genes were significantly up-regulated and more than 100 genes were significantly down-regulated in NASH-associated liver fibrosis (Fig. 1B). More than 300 genes were significantly upregulated and more than 100 genes were significantly downregulated in AH-associated liver fibrosis (Fig. 1C). Further, an intersection analysis of the differentially expressed (up-regulated) genes identified from liver tissue samples in the three groups of patients mentioned above, and a total of 25 commonly up-regulated genes in HBV infection, NASH, and AH-associated liver fibrosis disease were identified. The PPI analysis revealed significant interactions between the majority of generally upregulated genes (Fig. 1D). GO functional analysis of commonly up-regulated genes revealed that these genes were highly enriched in the GO entries of collagen-containing extracellular matrix and extracellular matrix structural constituent processes (Fig. 1E). Analyses of KEGG pathway enrichment showed a substantial enrichment of genes primarily in the bile secretion and focal adhesion pathways (Fig. 1F).

**FIG.1.**
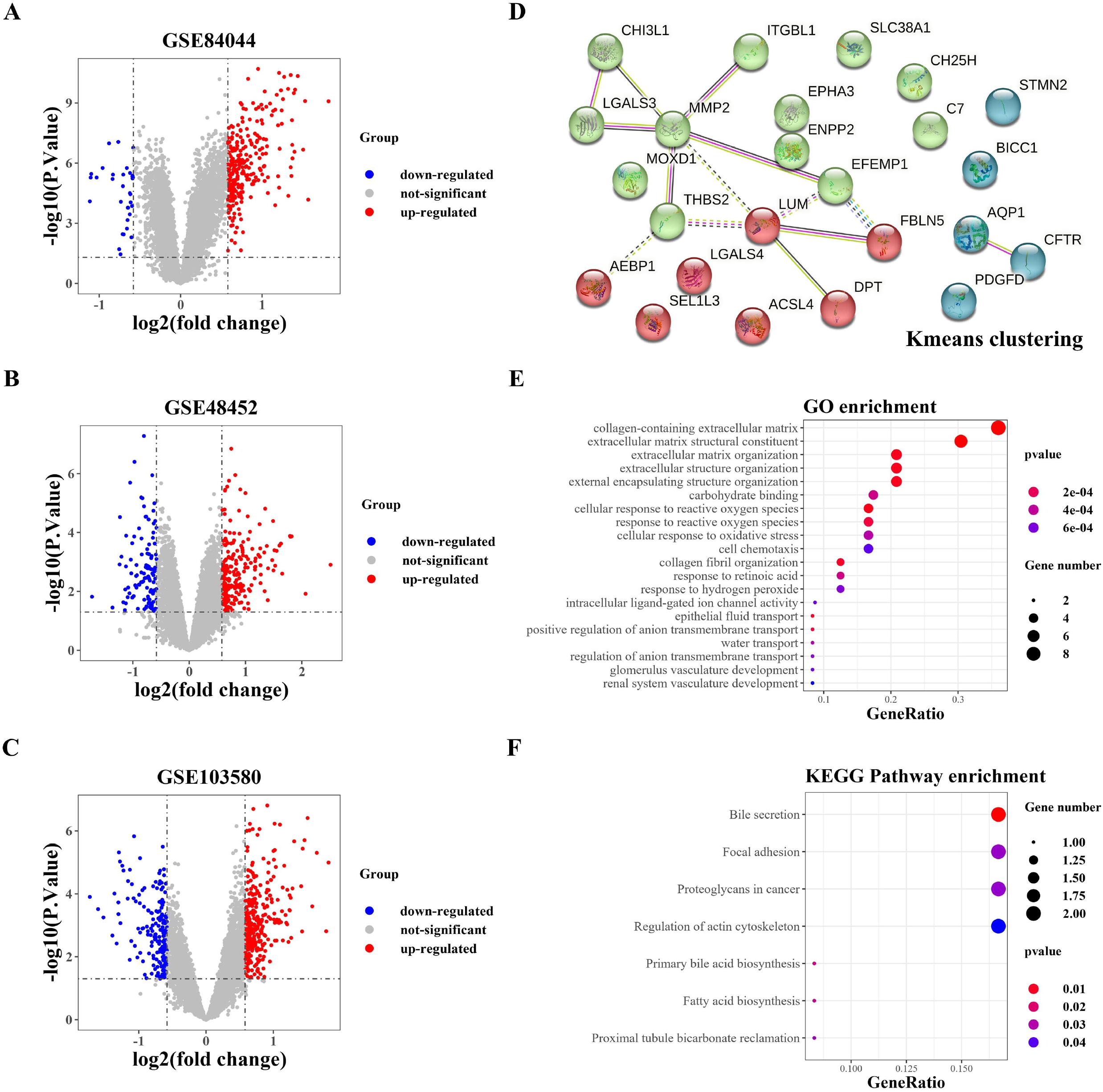
Screening and functional analysis of differentially expressed (upregulated) genes common to liver fibrosis induced by different etiologies. (A) Volcano map illustrating the differentially expressed genes in patients with HBV infection-associated liver fibrosis (GSE84044). (B) Volcano map of differentially expressed genes in patients with NASH-associated liver fibrosis (GSE48452). (C) Volcano map of differentially expressed genes from the dataset of patients with AH-associated liver fibrosis (GSE103580). (D) PPI analysis of commonly up-regulated genes in String database. (E) GO functional analysis of 25 commonly up-regulated genes. (F) KEGG pathway enrichment analysis of 25 commonly up-regulated genes.

### Ferroptosis-related genes were participated in liver fibrosis

Ferroptosis, a new cell death pattern, has an important biological function in several diseases. In this study, we obtained ferroptosis-related genes (promoter, suppressor, and biomarker, etc.) from FerrDb website and took intersection with 25 commonly up-regulated genes and found *ACSL4* and *SLC38A1* as ferroptosis-related drivers and *ENPP2* as ferroptosis-related suppressor. Further analysis of the relevant research literature demonstrated that a total of 10 genes were associated with ferroptosis, to the best of our knowledge,namely *AEBP1, THBS2, LGALS3, AQP1, EFEMP1, MMP2, ACSL4, SLC38A1, ENPP2* and *CFTR*.

Carbon tetrachloride (CCl_4_) is a hepatotoxin that can cause inflammation, damage, hepatocyte steatosis and liver fibrosis. CCl_4_-induced liver fibrosis mice model is an experimental animal model of liver fibrosis with more applications in basic research and drug evaluation because of its simple modeling, short cycle and reflecting the mechanism of liver fibrosis^[35-37]^. We screened three CCl_4_-induced liver fibrosis mice model datasets (GSE55747, GSE130123 and GSE152250) in GEO database and carried out expression analysis of 10 ferroptosis-related gene, and plotted heatmaps and box plots. We showed that *Acsl4* was significantly differentially expressed in the liver tissue dataset of three groups of CCl_4_-induced liver fibrosis mice model was significant differential. In addition, genes that showed significant differences in all three data sets included *Mmp2, Aebp1, Lgals3*, and *Thbs2*, while the expression levels of genes such as *Slc38a1, Aqp1, Efemp1, Enpp2*, and *Cftr* were not significantly different in all CCl_4_-induced liver fibrosis mice model datasets (Fig. 2).

**FIG.2.**
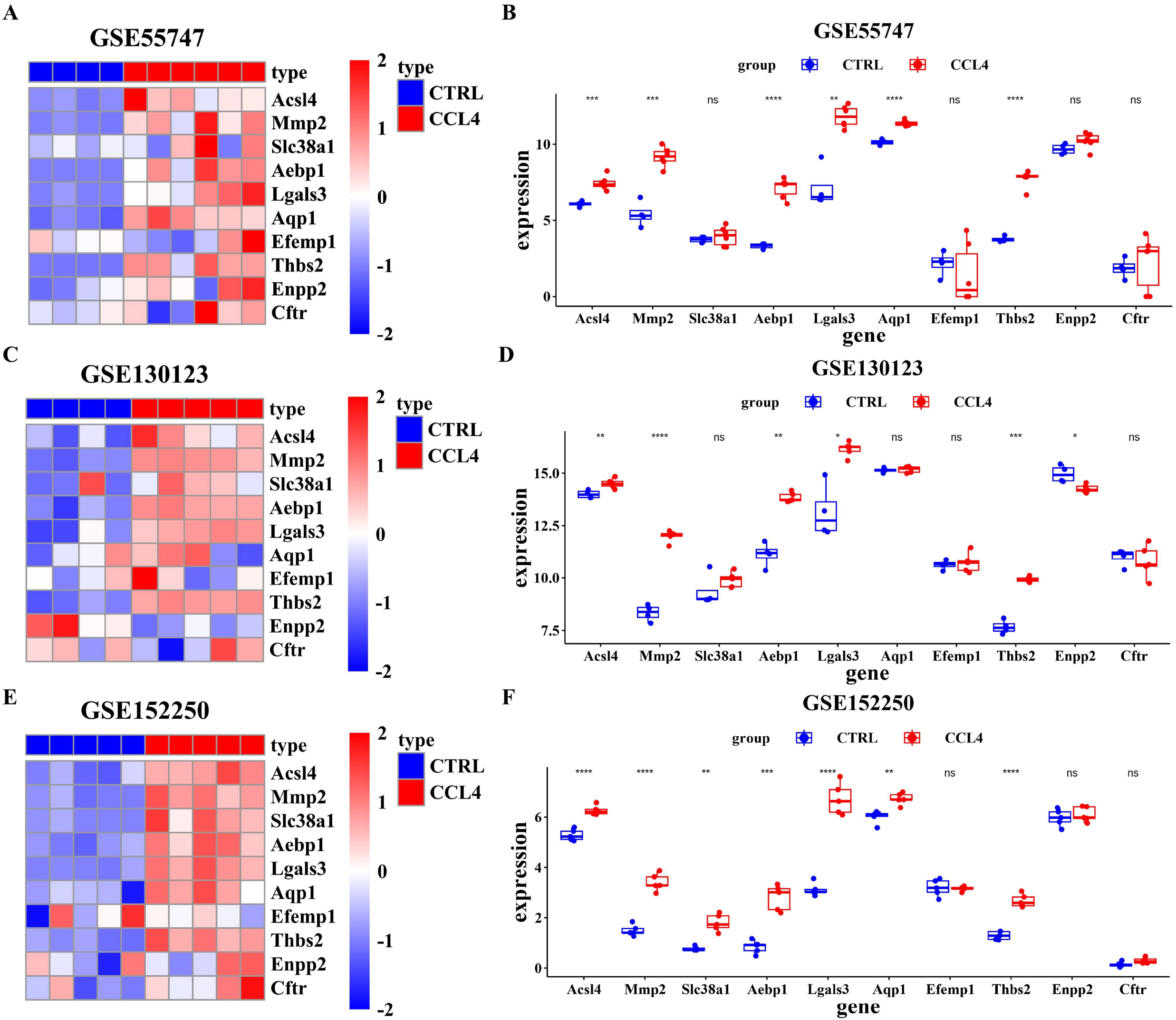
Validation of commonly up-regulated genes in a mice model of liver fibrosis. (A) Heatmap showing 10 Ferroptosis-Related Genes expression in GSE55747 dataset. (B) Box plot of 10 Ferroptosis-Related Genes expression in GSE55747 dataset. (C) Heatmap of 10 Ferroptosis-Related Genes expression in GSE130123 dataset. (D) Box plot showing the significance of 10 Ferroptosis-Related Genes expression in GSE130123 dataset. (E) Heatmap representing the significance of 10 Ferroptosis-Related Genes expression in GSE152250 dataset. (F) Box plot of the significance of 10 Ferroptosis-Related Genes expression in GSE152250 dataset.

### Transcriptomic analysis of liver fibrosis at various stages highlights the diagnostic importance of *ACSL4* in distinguishing severe cases

To determine whether the above 10 ferroptosis-related genes can distinguish severe liver fibrosis, we grouped samples from HBV infection-associated liver fibrosis patients (GSE84044), NASH-associated liver fibrosis patients (GSE48452), AH-associated liver fibrosis patients (GES103580), and HCC patients (GSE14520), as a way to study the different gene expression profile in severe liver fibrosis patients.

HBV infection-associated liver fibrosis patients were divided into three groups: no fibrosis (S0), early stage fibrosis (S1-2) and advanced stage fibrosis (S3-4) ^[38]^, then a multi-group analysis (Fig 3A) was performed for 10 genes, and the results revealed that *ACSL4* was specifically upregulated in liver tissues of HBV patients at advanced stage fibrosis (S3-4). NASH-associated liver fibrosis patients with control samples were divided into three groups: no fibrosis (F0), early-stage fibrosis (F1-2) and advanced stage fibrosis (F3-4), and then the genes were also analyzed for significant expression differences among multiple groups (Fig 3B), and seven genes, including *ACSL4, AEBP1, THBS2, LGALS3, AQP1, EFEMP1* and *CFTR*, were found to be specifically expressed in the period of advanced stage fibrosis (F3-4). AH-associated liver fibrosis patients were divided into three groups: mild acute alcoholic hepatitis, alcoholic steatosis and alcoholic cirrhosis (Fig 3C), and the results showed that all 10 genes were specifically expressed in alcoholic cirrhosis. The HCC patients’ adjacent non-tumor liver tissues were divided in two groups without and with cirrhosis, and revealed that nine genes, except *ENPP2*, were differential up-regulation during cirrhosis (Fig 3D). Among them, *ACSL4* showed specificity in severe liver fibrosis for different etiologies of patients. Further, gene expression time trend analysis was performed (Figure 4). Results showed that *ACSL4* appeared in the severe disease specific gene cluster with high specificity in all three data sets (Figure 4).

**FIG. 3.**
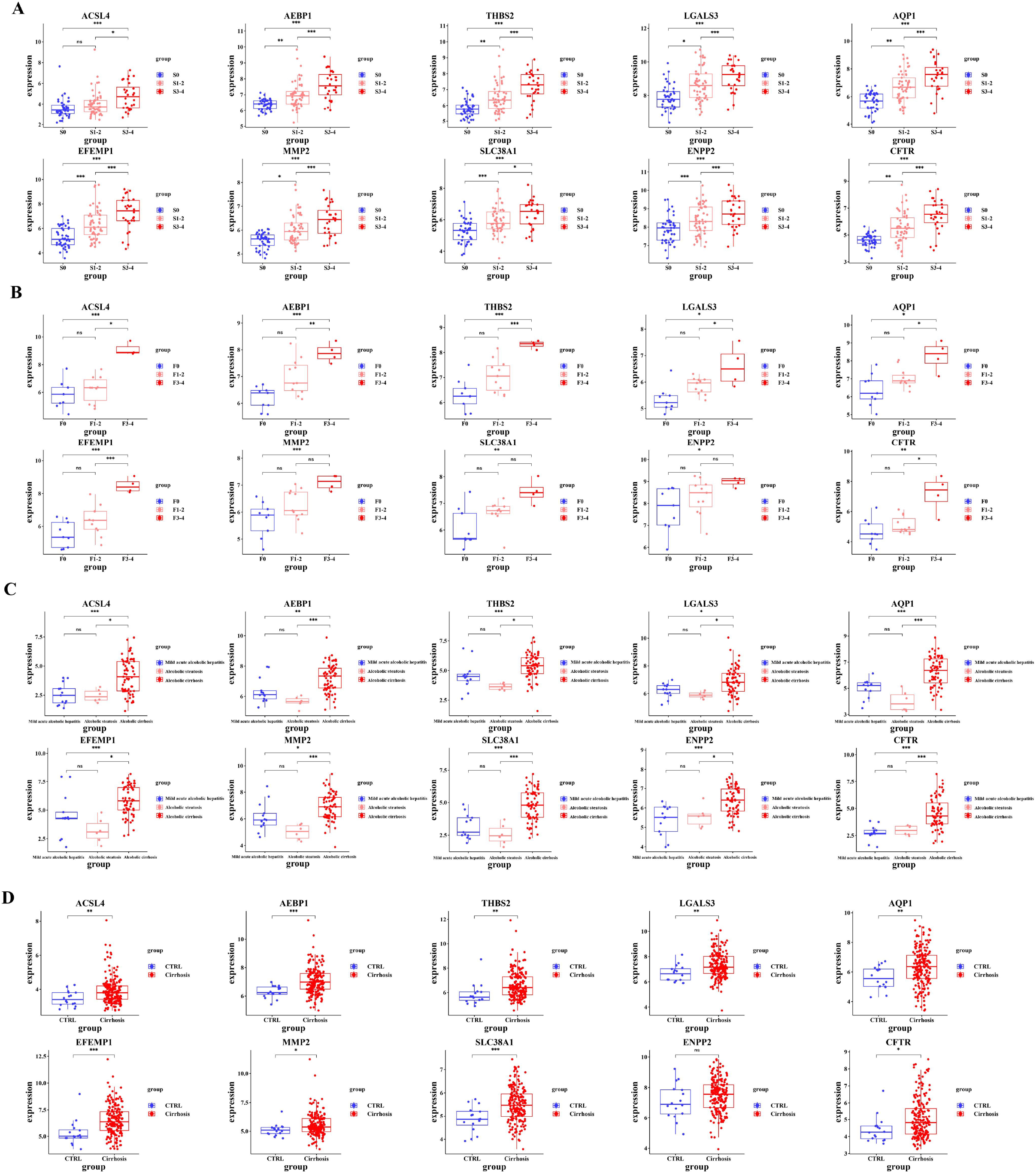
Analysis of clinical diagnostic value of *ACSL4* and nine other genes in patients with liver fibrosis. (A) Box plot illustrating the significance of 10 Ferroptosis-Related Genes expression in liver tissues of patients with HBV infection-associated liver fibrosis. (B) Box plot showing the significance of 10 Ferroptosis-Related Genes expression in liver tissues of patients with NASH-associated liver fibrosis. (C) Box plots of differential expression significance analysis of 10 Ferroptosis-Related Genes expression in liver tissues of patients with AH-associated liver fibrosis. (D) Box plot of differential expression significance analysis of 10 Ferroptosis-Related Genes expression in non-tumor tissue adjacent to the HCC.

**FIG. 4.**
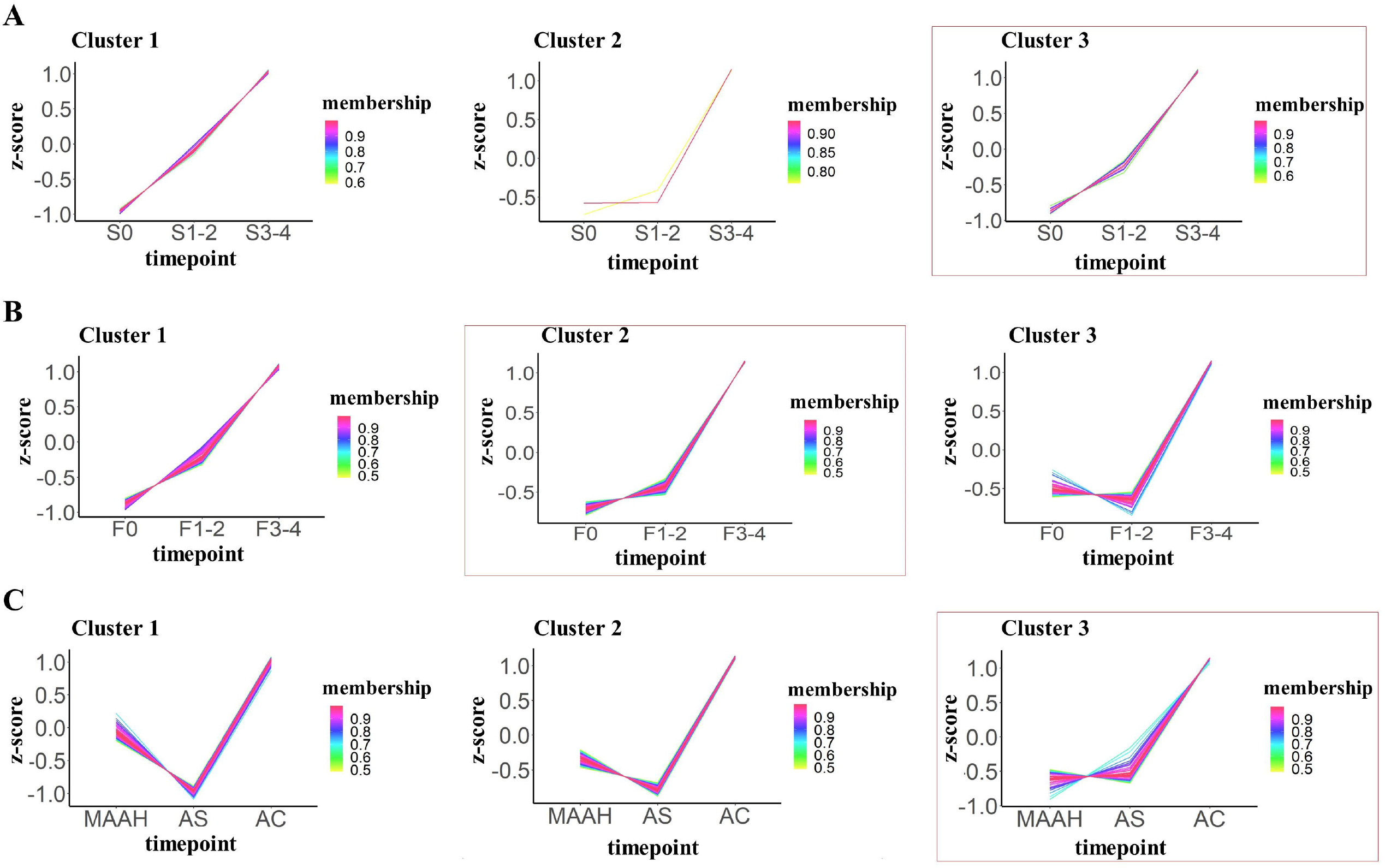
Gene expression time trend analysis of genes specific to HBV infection, NASH and AH-associated advanced liver fibrosis. (A) Clusters of expression time trends of differentially expressed (up-regulated) genes in the period of HBV 3). (B) Clusters of expression time infection-associated advanced hepatic fibrosis (S≥ trends of differentially expressed (up-regulated) genes in NASH infection-associated severe liver fibrosis (F≥3). (C) Clusters of expression time trends of differentially expressed (up-regulated) genes during AH infection-related liver fibrosis (alcoholic cirrhosis).

### Experimental validation confirmed the upregulation of *ACSL4* in advanced liver fibrosis induced by CCl_4_

Subsequently, our study was designed to experimentally verify the alterations in *Acsl4* expression levels by establishing a CCl_4_-induced mice model of advanced liver fibrosis. At the first step, mice were given an intraperitoneal injection of CCl_4_ twice a week for an 8-week period. Afterwards, the liver morphology and condition were examined upon sacrificing the mice. We observed the outcomes of H&E and Sirius Red staining of liver tissue samples The results revealed evident advanced liver fibrosis features in the liver tissues of mice in CCl_4_ group compared with control group. These features included disorganized liver lobule structure, disruption of normal hepatocyte architecture, inflammatory cells infiltration, fatty degeneration of liver, and significant fibrous connective tissue proliferation in the affected area (Fig 5A). Using the qPCR technique, the expression of *Acsl4* mRNA in mice liver tissues was evaluated (Fig. 5B). According to the findings, mice in the CCl_4_ group had significantly higher levels of *Acsl4* mRNA expression in their liver tissues than control mice.

**Fig. 5.**
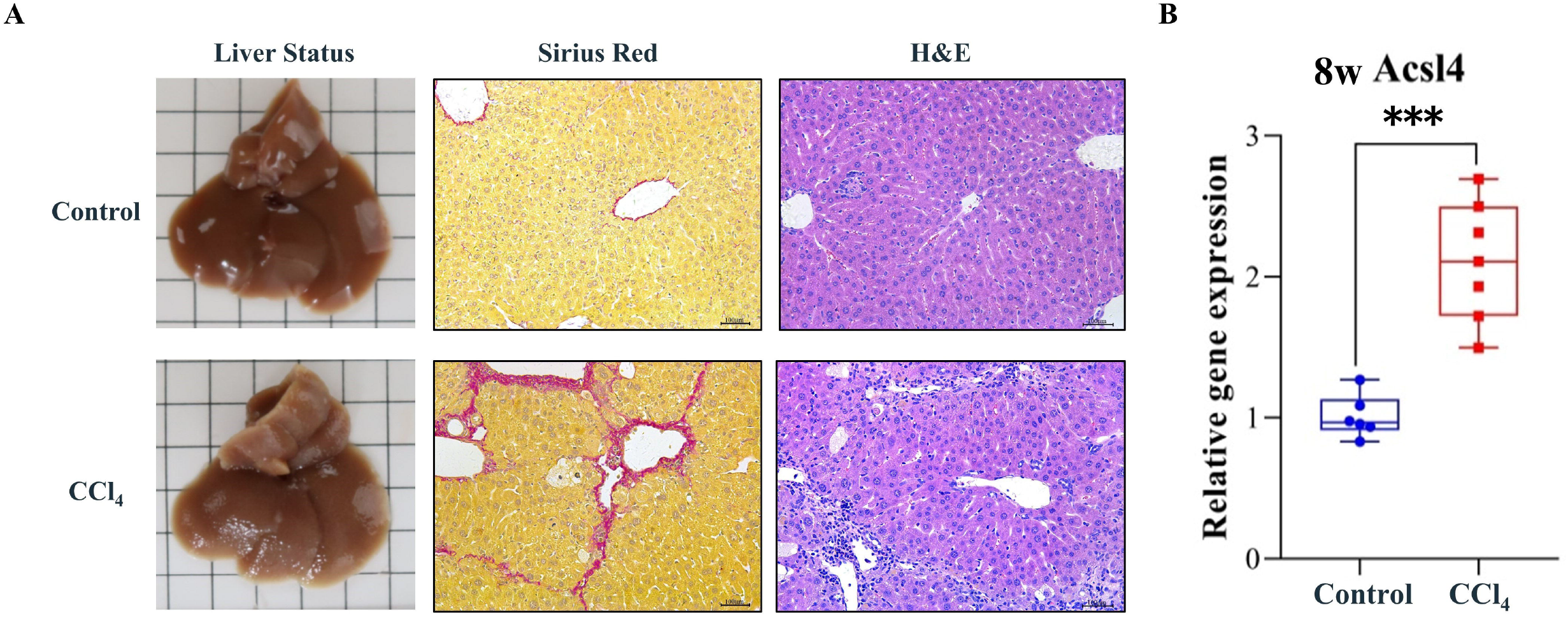
Mice model demonstration and validation of *Acsl4* gene expression levels. (A) Gross liver morphology of mice (left), Sirius red staining of liver tissue (middle) and H&E staining (right) in 1 mL/kg modeled 8-week control mice and CCl_4_ group mice. (B) The results of mRNA expression levels of *Acsl4* in liver tissues of control mice and CCl_4_ mice at 1 mL/kg for 8 weeks (n=6 for control group and n=7 for CCl_4_ group).

### *ACSL4* is intimately linked to crucial pathway component genes of liver fibrosis

The primary contributing factor to liver fibrosis and even cirrhosis is HBV infection, which has significantly raised the burden on healthcare in China and the Asia-Pacific area. We therefore extracted the TOP3 pathway and pathway component genes from the enrichment analysis of GO term and KEGG pathways of co-regulated genes in FIG 3 to give a preliminary understanding of the mechanisms by which *ACSL4* might exert a regulatory role in HBV-associated liver fibrosis and affect the development and progression of liver fibrosis. The correlation coefficient analysis of *ACSL4* with important pathway constitutive genes was performed using the HBV infection-associated liver fibrosis patient dataset (GSE84044), which includes a large number of liver fibrosis cases at different stages. The results revealed significant correlations between *ACSL4* and most of the component genes, including 12 genes such as *LUM, THBS2, EFEMP1*, and *CFTR* (Figure 6A∼L). Among them, we found that ACSL4 was significantly correlated with LUM in the constituent genes of the GO term (collagen-containing extracellular matrix, extracellular matrix structure and extracellular matrix organization) of TOP3 (Figure 6M∼O), and *CTRF, THBS2* and LUM in the constituent genes of the KEGG pathway (bile secretion, focal adhesion and proteoglycans in cancer) of TOP3, respectively correlation (Figure 6P∼R).

**FIG. 6.**
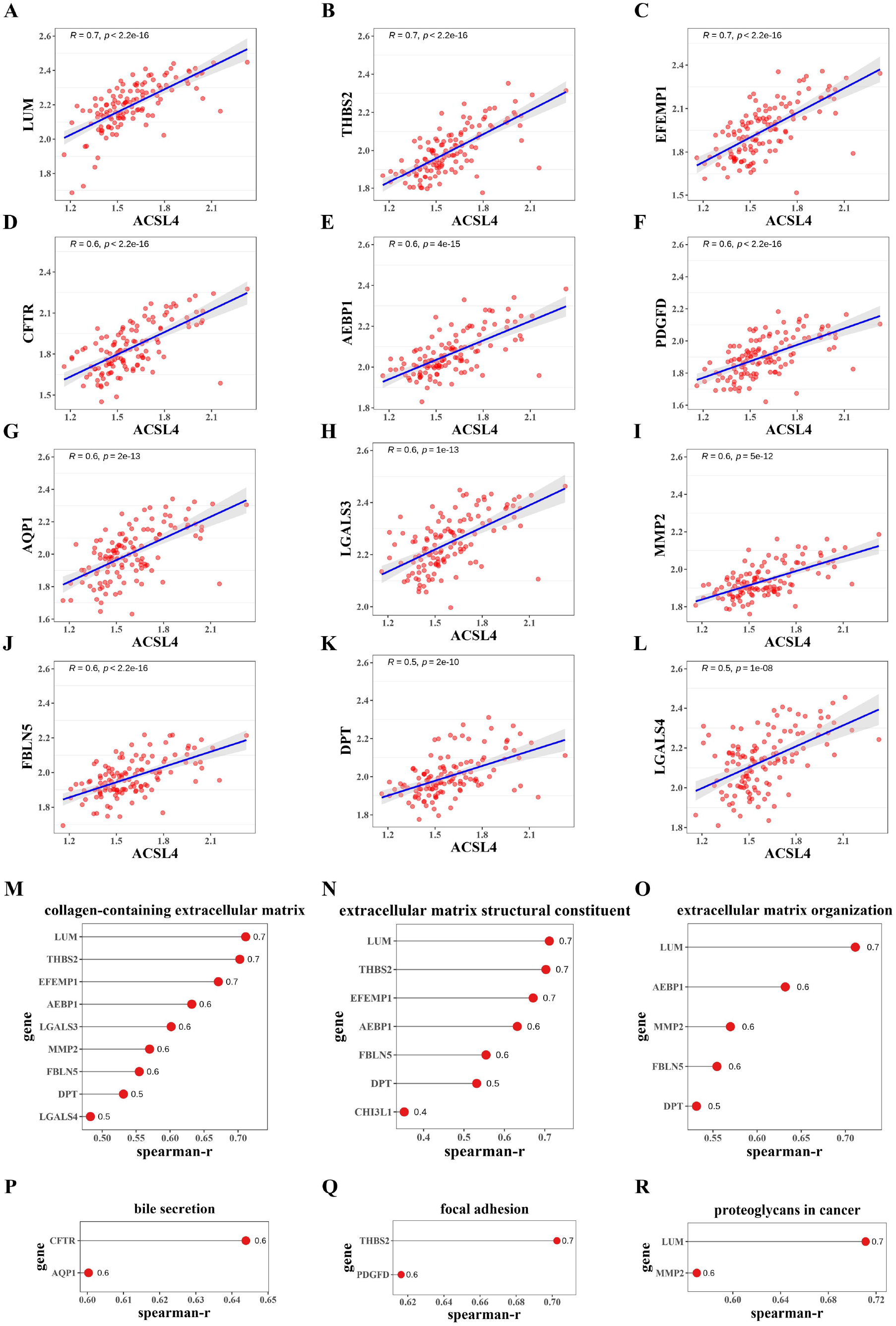
Correlation analysis of *ACSL4* with constitutive genes involved in liver fibrosis-related GO and KEGG pathways. (A∼L) Correlation analysis of *ACSL4* with 12 constitutive genes obtained from GO and KEGG pathway enrichment analyses. (M) Lollipop chart showing the correlation analysis between *ACSL4* and 9 constitutive genes involved in GO term: collagen-containing extracellular matrix. (N) Lollipop chart showing the correlation analysis between *ACSL4* with 7 constitutive genes involved in the GO term: extracellular matrix structural. (O) Lollipop chart showing the correlation analysis between *ACSL4* and 5 constitutive genes involved in GO term: extracellular matrix organization. (P) Correlation analysis of *ACSL4* with genes involved in the KEGG pathway: bile secretion. (Q) Correlation analysis of *ACSL4* with genes involved in the KEGG pathway: focal adhesion. (R) Correlation analysis of *ACSL4* with genes involved in the KEGG pathway: proteoglycans in cancer.

## Discussion

At present, a large number of available gene expression data (gene chip, RNA-Sequencing) have provided sufficient bioinformatics analysis conditions for this study. To identify genes with common and differential expression in the liver tissues of HBV infection-related liver fibrosis patients, NASH-related liver fibrosis patients, and AH-related liver fibrosis/cirrhosis patients, we collected multiple data sets and carried out data screening taking sample characteristics into consideration^[39]^.

Recently, in several bioinformatics-based studies on liver fibrosis, the identification of key genes related to the condition and the prediction of disease severity have primarily been limited to a single pathogenic factor. In 2021, a transcriptomics analysis identified up-regulated *THBS2* in NASH and advanced fibrosis (F3∼F4)^[40]^ and assessed its diagnostic capabilities by ROC curves. In 2022, another study demonstrated that the combined biomarkers of *ADAMTSL2* protein and eight proteins (*NFASC, CHST11, COLEC2, POR, FAH, SELE, THBS2*, and *TREM1*) were associated with the risk for and degree of fibrosis in NASH by multiplex analysis of patients with NAFLD in serum samples^[41]^, and pathological staging studies were also conducted using ROC curve analysis. The majority of studies for biomarker evaluation have focused on NASH, with HBV infection-related liver fibrosis being less frequently investigated. Although these studies have been successful in the diagnosis of liver fibrosis associated with NASH^[42]^, most of them only consider a single type of liver fibrosis, and it is difficult to expand its application to liver fibrosis diseases mediated by other causes. In this study, we combined bioinformatics analysis and literature research to identify ferroptosis-related genes such as *ACSL4, AEBP1, THBS2, LGALS3, AQP1, EFEMP1, MMP2, SLC38A1, ENPP2* and *CFTR*, which are aberrantly expressed in HBV infection, NASH and AH-associated liver fibrosis diseases, and it is anticipated that drugs targeting these ferroptosis-related genes will be used to treat various liver fibrosis illnesses. Literature research and analysis results showed that *ACSL4* is a key player in the ferroptosis pathway, and *ACSL4* has been proved in other models to lead to the ferroptosis cascade after the loss of *GXP4* activity, and inhibiting *ACSL4* may prevent ferroptosis^[28]^. Consistently, this study also revealed that *ACSL4* existed in the fatty acid biosynthesis in the enrichment analysis of KEGG pathway.

Liver fibrosis represents an aforementioned dynamic and treatable disease process^[43, 44]^, which is reversed to varying degrees in both patients and animals with lliver fibrosis once the cause is removed ^[45]^. Recent research results have shown that *Acsl4* has a trend of differential expression (up-regulation) in the NASH-related mice liver fibrosis model liver tissue, and has a close relationship with the progression of liver fibrosis in NASH-associated mice ^[33]^. However, *Acsl4* in the CCl_4_-induced liver fibrosis mice model and its biological function have not been full reported yet. In this study, we discovered that mice with severe liver fibrosis caused by CCl_4_ have much higher levels of *Acsl4* in liver tissue. Further exploration of the effectiveness of *Acsl4* server liver fibrosis, and in-depth explore the related functions of *Acsl4* to understand the pathogenic mechanism of *Acsl4*-related server liver fibrosis, which is promising to offer novel ideas and approaches for subsequent targeted drug therapy.

Liver biopsy as the “gold standard” for the diagnosis of liver fibrosis^[46]^, is generally performed with routine staining after serial sections of liver puncture samples and the addition of immunohistochemical staining for protein molecules according to the disease type, etc. It can also be combined with detection techniques such as PCR and enzymology^[47]^. However, the histological evaluation error and observer error existing in liver biopsy can limit the diagnosis and staging of fibrosis ^[48]^. In addition, the diagnostic biomarkers applied clinically still lack specificity and sensitivity to a certain extent to accurately diagnose the phase of liver fibrosis^[49]^. Bioinformatics methods can help to accurately identify clinical biomarkers of liver fibrosis, and relevant research has shown that VCAM-1 is a biomarker for the diagnosis of alcoholic cirrhosis^[50]^. *CXCL8, CXCL9, CXCL10* and *CXCL11* genes can be used as biomarkers of HBV infection-related liver injury^[51]^. In addition, gene mutations such as *ABCB4, ALDOB, GBE1, FAH, ASL, SLC25A13*, and *SERPINA1* have been shown at the cellular level to be closely associated with liver fibrosis ^[52]^. In this study, we found that *ACSL4* mRNA is preferentially enriched in liver fibrosis severe stage and has the ability to distinguish between early stage liver fibrosis patients and severe stage liver fibrosis patients through multiple comparative analysis of gene expression between groups and gene expression time trend analysis.

In summary, the study identified and obtained the upregulated ferroptosis-related gene *ACSL4*, which is consistently observed transcriptomic enriched in patients with HBV infection, NASH, AH-associated server liver fibrosis and cirrhotic tissue adjacent to hepatocellular carcinoma. Significant upregulation of *Acsl4* was verified in CCl_4_-induced advanced liver fibrosis mice model. Correlation analysis revealed that *ACSL4* was strongly correlated with *LUM, THBS2* and other 10 genes that constitute liver fibrosis key pathways. In summary, we highlight the preferential enrichment of *ACSL4* in severe liver fibrosis patients.

Our study has some limitations, including the need for a more extensive analysis of clinical samples and a broader inclusion of liver fibrosis etiologies. Future research should take experimental validation of hepatic *ACSL4* expression in patients into consideration. In addition to the significant utility of hepatic *ACSL4* as a clinical biomarker, further investigation is necessary to address its biological functions in advanced liver fibrosis.

